# Robust SARS-CoV-2 Neutralizing Antibodies Sustained through Three Months Post XBB.1.5 mRNA Vaccine Booster

**DOI:** 10.1101/2024.02.16.580687

**Authors:** Qian Wang, Ian A. Mellis, Yicheng Guo, Carmen Gherasim, Riccardo Valdez, Aubree Gordon, Lihong Liu, David D. Ho

## Abstract

SARS-CoV-2-neutralizing antibodies were substantially expanded one month after a shot of XBB.1.5 monovalent mRNA vaccine (XBB.1.5 MV) booster, but the durability of this response remained unknown. Here, we addressed this question by performing neutralization assays on four viral variants (D614G, BA.5, XBB.1.5, and JN.1) using sera from 39 adult participants obtained at ∼1 month and ∼3 months post an XBB.1.5 MV booster. Our findings indicate that the resultant neutralizing antibody titers were robust and generally maintained at stable levels for the study period, similar to those following XBB infection. Importantly, this durability of neutralizing antibody titers contrasts with the decline observed after a booster of the original monovalent or BA.5 bivalent mRNA vaccine. Our results are in line with the recent national data from the Centers for Disease Control and Prevention, showing the efficacy against symptomatic SARS-CoV-2 infection is sustained for up to 4 months after an XBB.1.5 MV booster.

## Main Text

In the Fall of 2023, the United States Food and Drug Administration authorized new COVID-19 vaccines to replace the previous bivalent vaccines^1^. The updated vaccines target the spike protein of SARS-CoV-2 Omicron subvariant XBB.1.5, aiming to enhance protection against the viral variant that was most dominant at the time strain selection was made^2^. Moreover, the updated vaccines were monovalent, because the prior ancestral/BA.5 bivalent vaccines did not appreciably expand the breadth of virus-neutralizing antibodies due to immunological imprinting that biased immune responses toward the ancestral strain^3-7^. We and others have since demonstrated that new updated XBB.1.5 mRNA monovalent vaccines (XBB.1.5 MV) boosted the potency and breadth of serum neutralizing antibodies against not only XBB.1.5 but also other Omicron subvariants, including the currently dominant JN.1, approximately one month after administration^2,8-15^. The removal of the ancestral spike from the new vaccine formulations had seemingly mitigated but not eliminated the immunological imprinting observed with the previous bivalent vaccines. However, how the serum antibody responses evolve in the ensuing months after an XBB.1.5 MV booster remains unknown.

We addressed this question by evaluating the serum virus-neutralizing titers at two distinct time points in 39 participants distributed across four cohorts: 1) who had received an XBB.1.5 MV booster without history of SARS-CoV-2 infection (XBB.1.5 MV); 2) who had XBB sublineage virus infection without history of XBB.1.5 MV booster (XBB infx); 3) who had received an XBB.1.5 MV booster with a prior pre-XBB Omicron infection history (pre-XBB Omicron infx + XBB.1.5 MV); and 4) who had received an XBB.1.5 MV booster with a prior XBB sublineage infection history (XBB infx + XBB.1.5 MV) (**Figure 1A**). The first and second serum samplings occurred at 26.8 and 82.1 mean days post vaccination or infection, respectively (**Figure 1A** and **Table S1**). No remarkable differences were noted in demographics or vaccination histories among the four clinical cohorts (**Tables S1 & S2**). Serum neutralizing antibody titers were determined using VSV-based pseudoviruses expressing the spike proteins of D614G (ancestral), BA.5, XBB.1.5, or JN.1.

**Figure 1.**
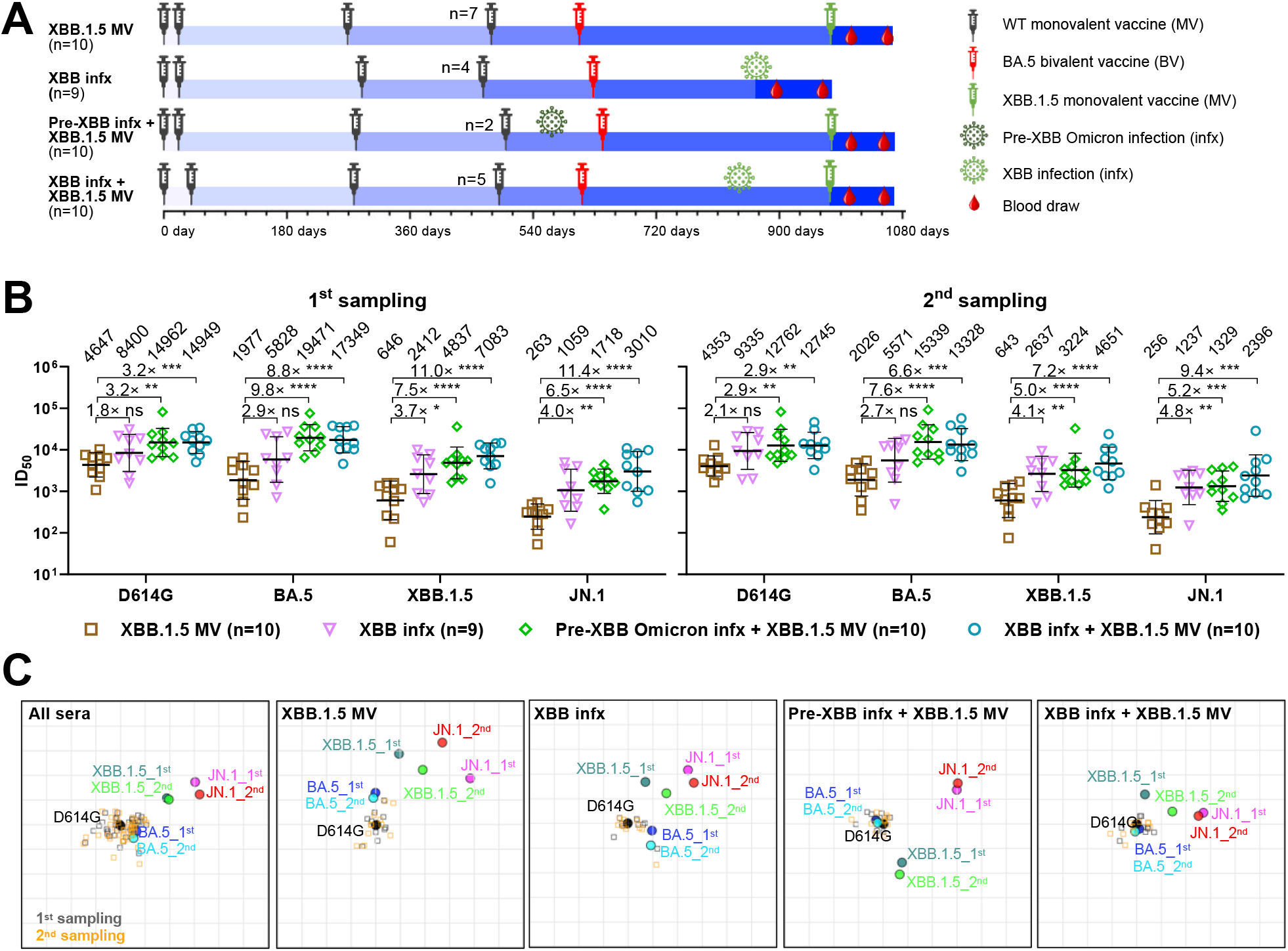
Neutralizing antibody titers in the months after an XBB.1.5 mRNA booster, XBB infection, or both. **A**. Timelines of vaccine administration, SARS-CoV-2 infection, and serum collection intervals for the four clinical cohorts in this study. Indicated time points represent the mean in days since first SARS-CoV-2 vaccination for each participant; day 0 is defined as the day of the initial SARS-CoV-2 vaccination. All participants previously received 3-4 doses of wildtype (WT) monovalent vaccines (MV), followed by one dose of the BA.5 bivalent vaccine (BV) booster. Numbers of participants for each group receiving a fourth WT MV is indicated. Serum samples were collected from two timepoints after an XBB.1.5 MV booster or XBB sublineage infection, as indicated, and the mean sample collection days post vaccination or infection are summarized in **Table S1**. n, sample size. **B**. Serum virus-neutralizing titers (ID50) of the four cohorts against the indicated SARS-CoV-2 pseudoviruses. Geometric mean ID50 titers (GMT) are shown along with the fold-differences in GMT versus the XBB.1.5 MV cohort. Statistical analyses comparing GMT between cohorts were performed by Mann-Whitney U tests. ns, not significant; \**p*<0.05; \*\**p*<0.01; \*\*\**p*<0.001; \*\*\*\**p*<0.001. **C**. Overlaid antigenic maps for serum virus-neutralizing titers at the first and second sampling timepoints. Panels in order: all cohorts, the XBB.1.5 monovalent vaccine (XBB.1.5 MV) cohort, the XBB infection (XBB infx) cohort, the Pre-XBB infection + XBB.1.5 monovalent vaccine (Pre-XBB infx + XBB.1.5 MV) cohort, and the XBB infection + XBB.1.5 monovalent vaccine (XBB infx + XBB.1.5 MV) cohort. Each panel contains two overlaid antigenic maps generated using sera from two time points post XBB exposure independently, with the D614G variant aligned. The x-y orientation of the component maps for either 1^st^ sampling or 2^nd^ sampling is free, as only the relative distances between the variants within a sampling and the respective sampling’s sera are compared. Distances between 1^st^ sampling and 2^nd^ sampling variant points are not directly compared. One grid square on the antigenic maps corresponds to one antigenic unit, representing an approximately 2-fold change in ID50 titer. Variant positions are indicated by circles, while serum positions are denoted by gray squares (1^st^ sampling) or orange squares (2^nd^ sampling). See also **Table S1, Table S2, and Figure S1**.

Overall, similar patterns in neutralizing antibody titers (ID50) were observed between serum samples from the first and second time points for all four cohorts (**Figure 1B**), with several features worthy of emphasis. First, neutralizing titers against all four viruses tested were robust (≥256) at both time points. Second, the highest neutralizing titers of the XBB.1.5 MV cohort was observed against D614G, while titers against XBB.1.5 were much lower (4,647 versus 646 and 4,535 versus 643 at first and second time points, respectively). This finding showed the persistence of immunological back-boosting, although not as severe as previously observed for the ancestral/BA.5 bivalent vaccine boost^3-7^. Third, as expected, sera of participants from “pre-XBB Omicron infx + XBB.1.5 MV” and “XBB infx + XBB.1.5 MV” cohorts (individuals who had one more exposure to Omicron spike) showed substantially higher neutralizing titers against all four viruses. Fourth, sera from the XBB infx cohort exhibited stronger neutralizing activity (1.8-4.8-fold) than sera from the XBB.1.5 MV cohort. Lastly, among the viruses tested, serum neutralizing titers were the lowest for JN.1 in all cohorts, indicating that it is the most antibody-evasive, in line with our previous report^2^.

Antigenic maps were then generated using serum neutralization data from the four cohorts, both collectively and individually (**Figure 1C**), to visually summarize our findings. These maps showed that the antigenic distances between the ancestral D614G virus and other tested SARS-CoV-2 Omicron subvariants at both time points were consistent across all cohorts. Infections with an XBB sublineage virus seemed to slightly outperform a single XBB.1.5 MV booster in reducing antigenic distance. Additionally, having a history of Omicron infection prior to XBB.1.5 MV booster enhanced the neutralization of the currently dominant JN.1 subvariant. In particular, exposure to XBB sublineage infection before the booster appeared to reduce the antigenic distance to JN.1 by approximately one antigenic unit more than a pre-XBB Omicron infection.

We next endeavored to better understand the change in virus-neutralizing titers between the first and second serum samples. The ID50 titers against four viruses for all four cohorts were plotted using the actual dates of serum collection post vaccination or infection (**Figure 2A**). No significant waning of neutralizing titers was observed for all four cohorts, except for a slight decrease in titer against XBB.1.5 in the pre-XBB infx + XBB.1.5 MV cohort (1.5-fold decrease). However, in our prior study^4^, we observed around a 2-fold or greater decrease in neutralizing titers against D614G, BA.5, and XBB.1.5 from ∼1 month to ∼3 month post the fourth dose of the wildtype monovalent vaccine (WT MV) booster or ancestral/BA.5 bivalent (BV) booster, while a BA.5 breakthrough infection led to stable serum neutralizing activity over the same interval (**Figure 2B**). Since the sample collection times across the two studies were slightly different (**Figure S1** and **Table S1**), the most appropriate comparison of antibody decline would be to determine the slope between the ID50 titers for the two time points. Comparing cohorts with vaccine boosters but without any history or laboratory evidence of prior SARS-CoV-2 infection, the median slope for all four viruses tested was nearly zero (flat) after an XBB.1.5 MV booster in the present study, whereas the median slopes were significantly more negative (decline, with median values ranging from -0.0040/day to -0.0063/day) after a WT MV or ancestral/BA.5 BV booster in a prior study (**Figure 2C**). On the other hand, there was no appreciable decline during the study period for serum neutralizing titers for individuals who had a BA.5 or XBB breakthrough infection in both studies (**Figure 2D**).

**Figure 2.**
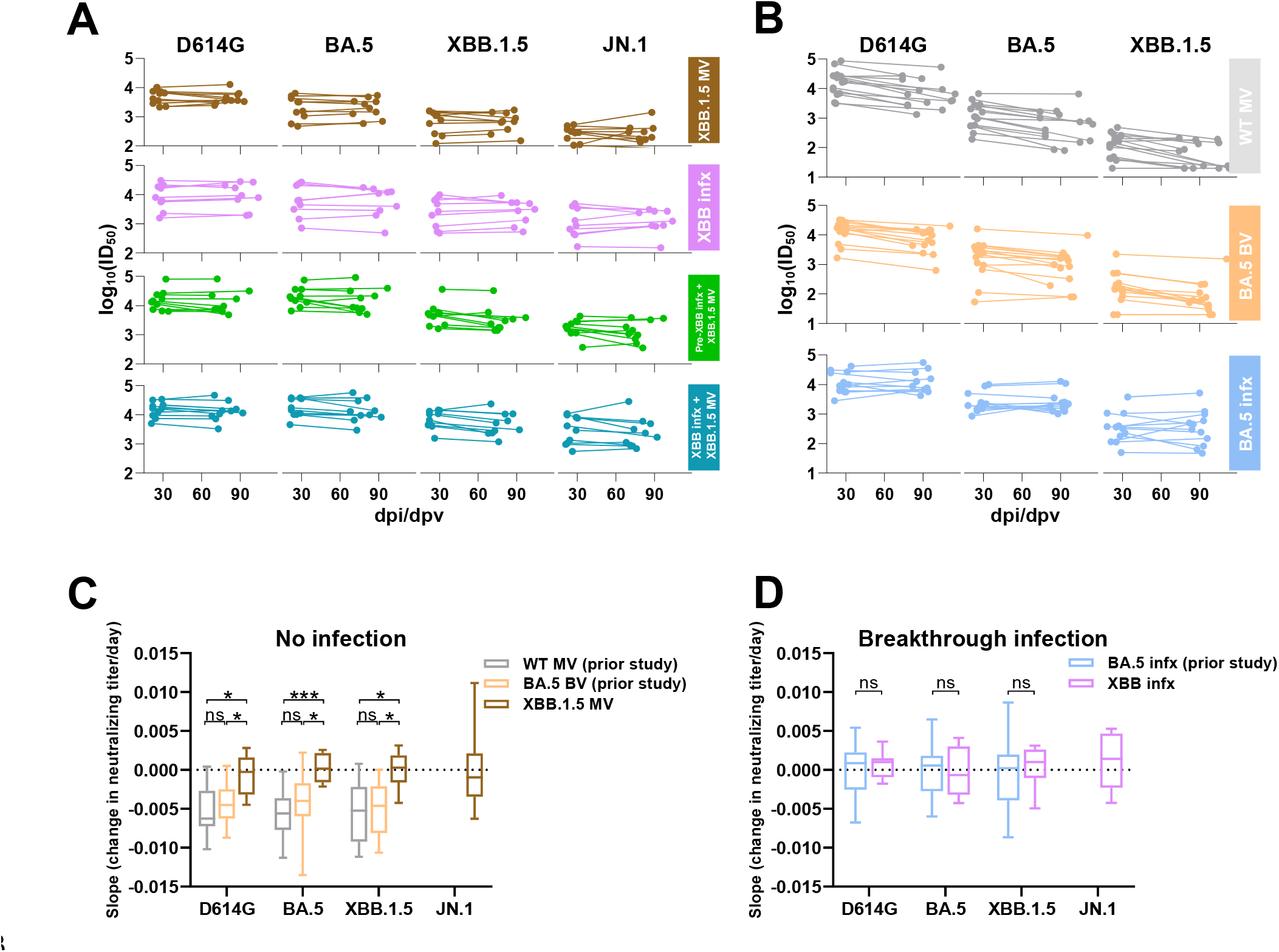
An XBB.1.5 monovalent vaccine booster induced sustained virus-neutralizing antibody responses through 3 months post vaccination. **A**. Comparison of serum neutralizing antibody titers between two timepoints after XBB vaccination or infection for each participant in each cohort. **B**. Comparison of serum neutralizing antibody titers between two timepoints after the 4^th^ dose of the WT MV or BA.5 BV, or BA.5 breakthrough infection. Data for “WT MV”, “BA.5 BV” and “BA.5 infx” cohorts were extracted from a previously published study^4^. **C-D**. Estimated change in neutralizing titer per day (i.e., slope from first to second titer) for the cohorts without SARS-CoV-2 infection (**C**) and with breakthrough infection (**D**) across the two respective studies. Negative values represent the speed of titer decay. Box-and-whisker plots, with whisker limits at minimum and maximum values and the central line representing median. Statistical analyses were performed by Mann-Whitney U tests. ns, not significant; \**p*<0.05; \*\*\**p*<0.001. See also **Figure S2**.

In summary, an XBB.1.5 MV booster has elicited robust and sustained neutralizing antibodies against SARS-CoV-2, including the currently dominant JN.1 subvariant, for approximately 3 months (**Figures 1 & 2**). The lack of waning of neutralizing antibodies is in distinct contrast to that observed after an ancestral MV or ancestral/BA.5 BV booster (**Figure 2C**). A clear explanation for this discrepancy is lacking. There is no indication that we may have missed the peak antibody response with the first serum sampling (**Figure S1**). While a SARS-CoV-2 infection between the two time points could not be excluded, antibodies directed to the viral nucleocapsid could not be detected in the sera from the XBB.1.5 MV cohort (data not shown), nor is there a clinical history consistent with an intervening infection. The short-term durability of the potency and breadth of the virus-neutralization response following XBB.1.5 MV booster again suggests that immunological imprinting has been partially mitigated, likely due to the exclusion of the ancestral spike from the updated vaccine formulation. However, it should be noted that immunological imprinting has not been completely overridden by two exposures to an Omicron spike, as evident by higher neutralizing antibody titers against D614G than against XBB.1.5 in the two cohorts that had an Omicron infection followed by an XBB.1.5 MV boost (**Figure 1B**). Importantly, our new data are in agreement with vaccine effectiveness estimates recently reported by the Center for Disease Control and Prevention, showing no appreciable decline in protection against symptomatic COVID-19 up to four months post booster with updated vaccines^16^.

## Limitations of the study

There are two notable limitations to this study. First, the number of study subjects in each cohort is relatively small, even though statistically significant results are reported. Second, we still lack an explanation for the mechanism by which neutralizing antibody titers after an XBB.1.5 MV booster is more stable than those elicited by earlier vaccinations. More studies are required to provide an answer.

## FIGURE AND LEGENDS

**Figure S1.**
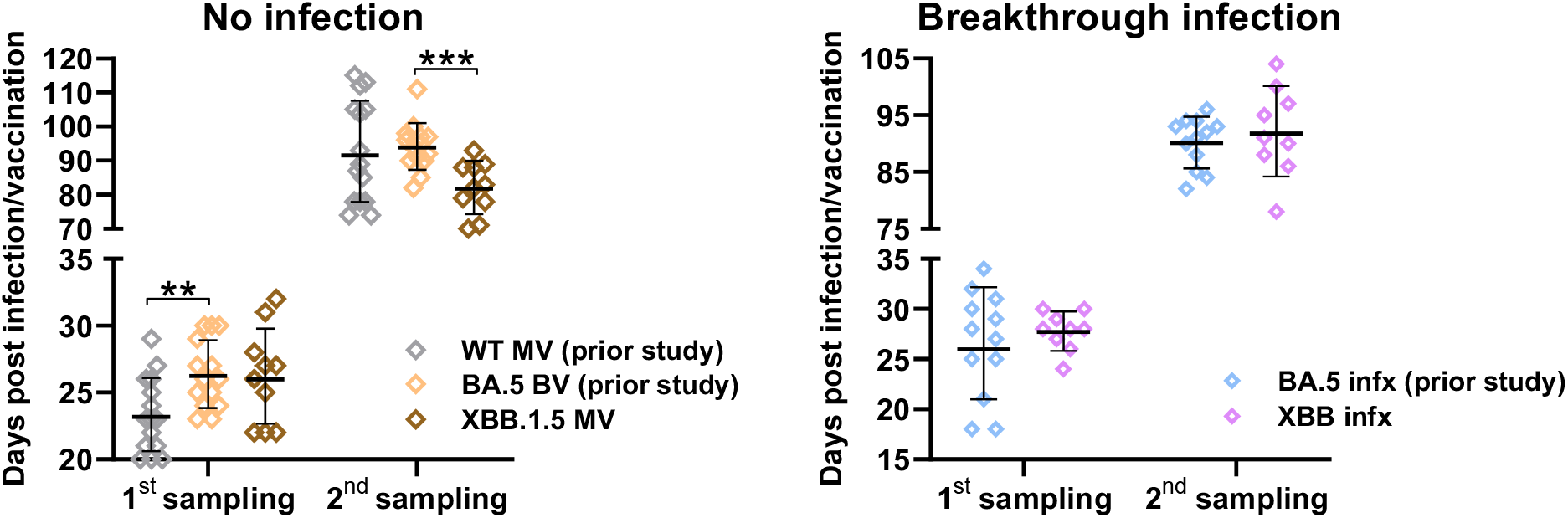
Comparison of sample collection days post-vaccination or infection, related to Figure 1. Statistical analyses were performed by Mann-Whitney U tests. \*\**p*<0.01; \*\*\**p*<0.001.

**Figure S2.**
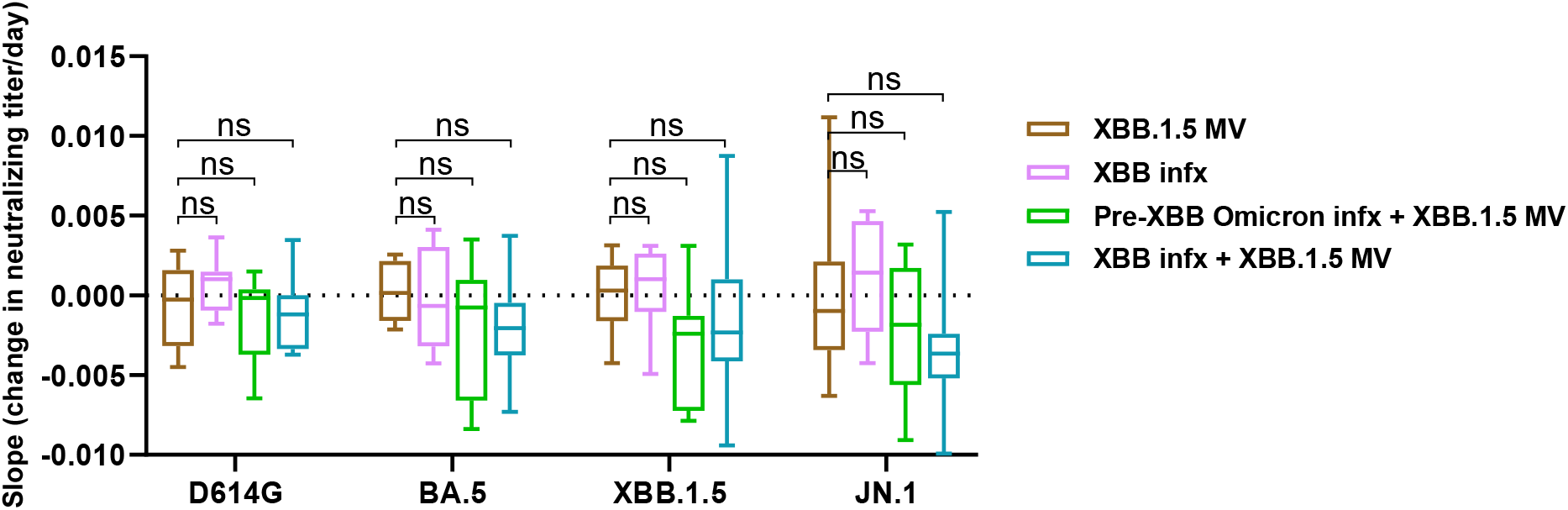
ID50 slope (change in neutralizing titer per day) for the indicated cohorts, related to Figure 2. Statistical analyses were performed by Mann-Whitney U tests. ns, not significant.

**Table S1.** Summary of participant information. Demographic, vaccine, and serum collection information summarized for each cohort. Listed values represent the mean and range (age and sera collection) or number and percentage (vaccine type and sex).

| Clinical Information |  | Across all |  | XBB.1.5 MV |  | XBB infx |  | Prior Omicron infection + XBB.1.5 MV |  |  |  |
| --- | --- | --- | --- | --- | --- | --- | --- | --- | --- | --- | --- |
|  |  |  |  |  |  |  |  | Pre-XBB infx + XBB.1.5 MV |  | XBB infx + XBB.1.5 MV |  |
|  |  | Number or mean | % or range | Number or mean | % or range | Number or mean | % or range | Number or mean | % or range | Number or mean | % or range |
| Total participants |  | 39 |  | 10 |  | 9 |  | 10 |  | 10 |  |
| Male |  | 8 | 20.5% | 3 | 30% | 1 | 11.10% | 2 | 20% | 2 | 20% |
| Female |  | 31 | 79.5% | 7 | 70% | 8 | 88.90% | 8 | 80% | 8 | 80% |
| Age |  | 51.6 | (30, 67) | 52.6 | (38, 65) | 52.1 | (40, 59) | 49.7 | (36, 67) | 52.1 | (30, 62) |
| First two WT doses | Pfizer | 35 | 89.7% | 9 | 90% | 8 | 88.90% | 9 | 90% | 9 | 90% |
|  | Moderna | 3 | 7.7% | 1 | 10% | 1 | 11.10% | 1 | 10% | 0 | 0% |
|  | Janssen | 1 | 2.6% | 0 | 0% | 0 | 0% | 0 | 0% | 1 | 10% |
| Third WT dose | Pfizer | 34 | 87.2% | 9 | 90% | 8 | 88.90% | 9 | 90% | 8 | 80% |
|  | Moderna | 5 | 12.8% | 1 | 10% | 1 | 11.10% | 1 | 10% | 2 | 20% |
| Fourth WT dose | Pfizer | 13 | 33.3% | 4 | 40% | 3 | 33.30% | 2 | 20% | 4 | 40% |
|  | Moderna | 5 | 12.8% | 3 | 30% | 1 | 11.10% | 0 | 0% | 1 | 10% |
|  | None | 21 | 53.8% | 3 | 30% | 5 | 55.60% | 8 | 80% | 5 | 50% |
| BA.5 bivalent booster | Pfizer | 23 | 59% | 6 | 60% | 7 | 77.80% | 4 | 40% | 6 | 60% |
|  | Moderna | 16 | 41% | 4 | 40% | 2 | 22.20% | 6 | 60% | 4 | 40% |
| XBB MV booster | Pfizer | 13 | 33.3% | 4 | 40% | 0 | 0% | 3 | 30% | 6 | 60% |
|  | Moderna | 17 | 43.6% | 6 | 60% | 0 | 0% | 7 | 70% | 4 | 40% |
|  | None | 9 | 23.1% | 0 | 0% | 9 | 100% | 0 | 0% | 0 | 0% |
| First sample post-XBB (days) |  | 26.4 | (21, 34) | 26.2 | (22, 32) | 27.8 | (24, 30) | 26.9 | (21, 34) | 24.7 | (21, 30) |
| Second sample post-XBB (days) |  | 82.1 | (68, 104) | 82.1 | (70, 93) | 92.1 | (78, 104) | 77.6 | (68, 97) | 77.5 | (68, 92) |

**Table S2.**
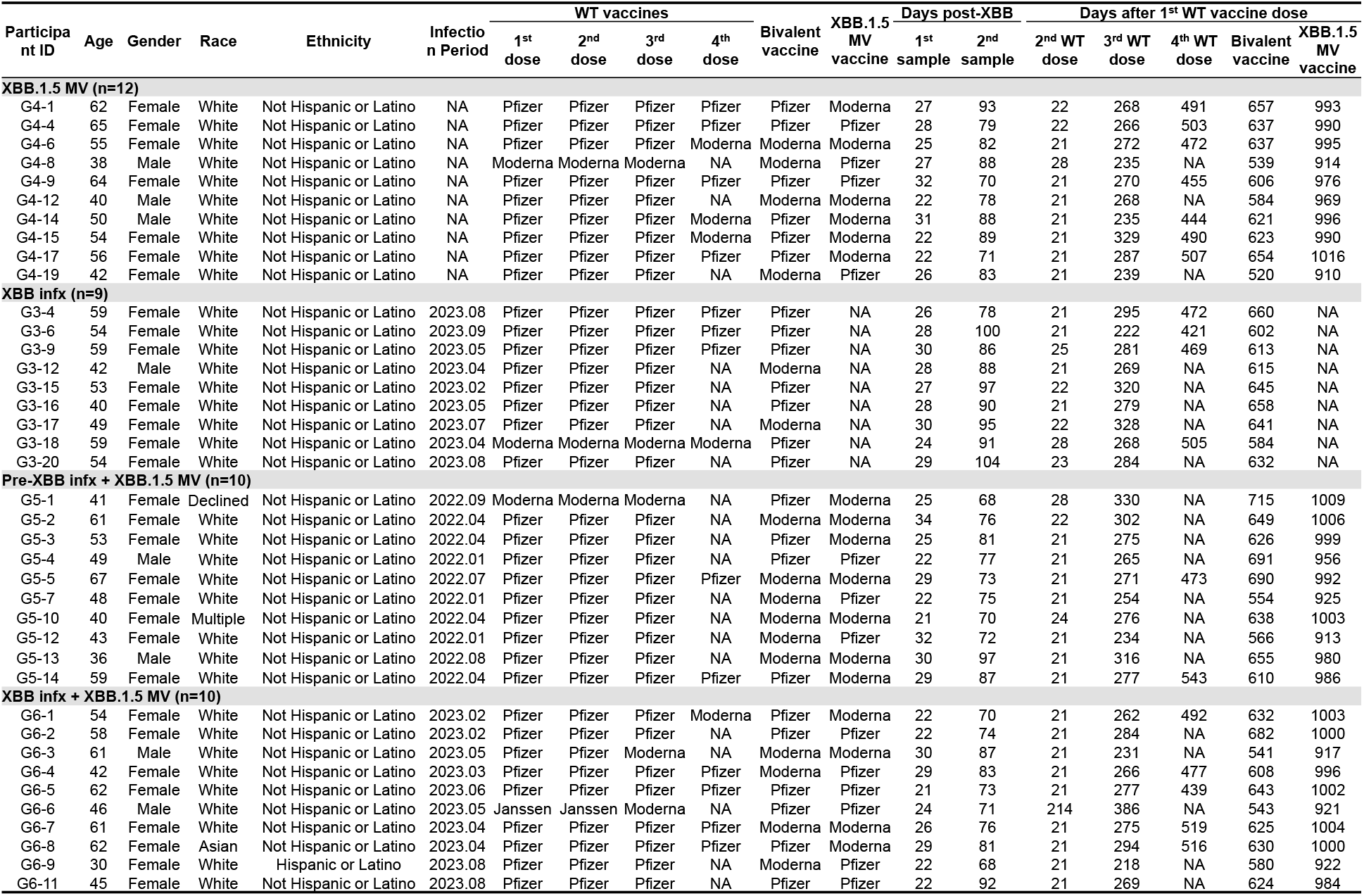
Participant details. Clinical information for each participant, including demographic, vaccine, infection, and sera details.

## STAR METHODS

### KEY RESOURCES TABLE

#### RESOURCE AVAILABILITY

∘ Lead contact
∘ Materials availability
∘ Data and code availability

### EXPERIMENTAL MODEL AND SUBJECT DETAILS

∘ Clinical cohorts
∘ Cell line

### METHOD DETAILS

∘ Pseudovirus neutralization assay
∘ Antigenic cartography

### QUANTIFICATION AND STATISTICAL ANALYSIS ACKNOWLEDGEMENTS

This study was supported by funding from the NIH SARS-CoV-2 Assessment of Viral Evolution (SAVE) Program (Subcontract No. 0258-A709-4609 under Federal Contract No. 75N93021C00014) and the Gates Foundation (project INV019355) to D.D.H., as well as funding from the NIH contract 75N93019C00051 to A.G. We express our gratitude to Zijin Chu, Theresa Kowalski-Dobson, Emily Stoneman, David Manthei, Anna Buswinka, Gabe Simjanovski, Joseph Wendzinski, Mayurika Patel, Kathleen Lindsey, Dawson Davis, Victoria Blanc, Savanna Sneeringer, and Pamela Bennett-Baker of the IASO study team for conducting the IASO study.

## AUTHOR CONTRIBUTIONS

L. L., A.G., and D.D.H. conceived and supervised this project. Q.W. managed the project. Q.W., I.A.M., and L. L. conducted pseudovirus neutralization assays. Y.G. and I.A.M. conducted bioinformatic analyses. C.G., R.V., and A.G. provided clinical samples. Q.W., I.A.M., Y.G., L. L., and D.D.H. analyzed the results and wrote the manuscript. All authors reviewed the results and approved the final version of the manuscript.

## RESOURCE AVAILABILITY

### Lead contact

Further information and requests for resources and reagents should be directed to and will be fulfilled by the Lead Contact Author David D. Ho.

### Materials availability

All reagents generated in this study are available from the Lead Contact Author with a completed Materials Transfer Agreement.

### Data and code availability

Data reported in this paper will be shared by the Lead Contact Author upon request. This paper does not report original code. Any additional information required to reanalyze the data reported in this paper and R scripts used to perform most statistical analyses are available from the Lead Contact Author upon request.

## EXPERIMENTAL MODEL AND SUBJECT DETAILS

### Clinical cohorts

Longitudinal sera were obtained as part of an ongoing cohort study, Immunity-Associated with SARS-CoV-2 Study (IASO), which began in 2020 at the University of Michigan in Ann Arbor, Michigan^17^. All participants provided written informed consent, and sera were collected according to the protocol approved by the Institutional Review Board of the University of Michigan Medical School. Participants in the IASO study completed weekly symptom surveys, and if any symptoms were reported, participants were tested for SARS-CoV-2. We tested all serum samples by anti-nucleoprotein (NP) ELISA to check for any potential breakthrough infections during the period spanning sample collections for this study.

In this study, we included sera from 39 individuals across four clinical cohorts: 1) individuals with no recorded SARS-CoV-2 infections who had received an XBB.1.5 monovalent vaccine booster (“XBB.1.5 MV”); 2) individuals with a recent XBB sublineage infection who had not received the XBB.1.5 booster (“XBB infx); 3) individuals with prior infection who also received the XBB.1.5 booster (“Pre-XBB Omicron infx + XBB MV”); and 4) individuals with an XBB sublineage infection who also received the XBB.1.5 booster (XBB infx + XBB MV). Individuals in all cohorts previously received either three or four doses of a wildtype monovalent vaccine as well as a single ancestral/BA.5 bivalent booster. Most participants were female (79.5%) with an average age of 51.6 years. Sera were collected an average of 26.4 and 82.1 days after XBB.1.5 vaccination or XBB sublineage infection. Demographic, vaccination, and serum collection details are summarized for each cohort in **Table S1**, and details are shown for each participant in **Table S2**.

### Cell lines

We obtained 293T (CRL-3216) and Vero-E6 (CRL-1586) cells from ATCC and cultured them according to the manufacturer’s instructions. The morphology of each cell line was visually confirmed before use. All cell lines tested negative for mycoplasma.

## METHOD DETAILS

### Pseudovirus neutralization assay

Plasmids encoding SARS-CoV-2 variant spikes, including D614G, BA.5, XBB.1.5, and JN.1, were generated in previous studies^2,5,18^.

To produce pseudotyped viruses of SARS-CoV-2 variants, we transfected 293T cells with the spike-encoding plasmids described above using 1 mg/mL PEI (Polyethylenimine). One day post-transfection, the 293T cells were incubated with VSVG*ΔG-luciferase (Kerafast, Inc.) at a multiplicity of infection of approximately 3 to 5 for 2 hours followed by three washes with complete culture medium. The cells were then cultured with fresh medium for an additional day. Cell supernatants containing viruses were collected, clarified by centrifugation, aliquoted, and stored at -80°C until use.

The viral titer of each variant was titrated to calculate a 50% tissue-culture-infectious dose (TCID50) and normalized for neutralization assays. Serum samples were diluted in triplicate in 96-well plates, starting from a 12.5-fold dilution (for one sample, a 50-fold starting dilution was necessary due to volume constraints), and then incubated with an equal volume of virus for 1 hour at 37°C before adding 2 × 10^4^ cells/well of Vero-E6 cells. The cells were then cultured overnight, harvested, and lysed for measurement of luciferase activity using SoftMax Pro v.7.0.2 (Molecular Devices). Reductions in luciferase activity at given dilutions of sera were calculated, and ID50 values of sera were obtained by fitting the virus-reduction data using a non-linear five-parameter dose-response curve in GraphPad Prism V.10.

### Antigenic cartography

Antigenic maps for D614G and other SARS-CoV-2 variants were generated by integrating the ID50 values of individual serum samples using a published antigenic cartography method^19^. Visualizations were created with the Racmacs package (version 1.1.4, available at https://acorg.github.io/Racmacs/) in R software version 4.0.3. The optimization was performed over 2,000 steps, with the ‘minimum column basis’ parameter set to ‘none’. The ‘mapDistances’ function calculated the antigenic distances between each serum sample and each variant. D614G was used as center for each group’s sera, and the seeds for each antigenic map were manually adjusted to ensure that XBB.1.5 and JN.1 were oriented correctly relative to D614G. All maps are aligned with D614G positioned in the middle left, ensuring a consistent reference point across the maps.

## QUANTIFICATION AND STATISTICAL ANALYSIS

Serum neutralization ID50 values were calculated using a five-parameter dose-response curve in GraphPad Prism v.10. Evaluations of statistical significance were performed employing Mann-Whitney U tests using GraphPad Prism v.10 software. Mann-Whitney U tests, unpaired t-tests, and Bonferroni correction were performed in R version 4.3.2.

## REFERENCES

1. FDA (2023). Updated COVID-19 Vaccines for Use in the United States Beginning in Fall 2023. https://www.fda.gov/vaccines-blood-biologics/updated-covid-19-vaccines-use-united-states-beginning-fall-2023.

2. Wang, Q., Guo, Y., Bowen, A., Mellis, I.A., Valdez, R., Gherasim, C., Gordon, A., Liu, L., and Ho, D.D. (2023). XBB.1.5 monovalent mRNA vaccine booster elicits robust neutralizing antibodies against emerging SARS-CoV-2 variants. bioRxiv, 2023.2011.2026.568730. 10.1101/2023.11.26.568730.

3. Wang, Q., Bowen, A., Valdez, R., Gherasim, C., Gordon, A., Liu, L., and Ho, D.D. (2023). Antibody Response to Omicron BA.4-BA.5 Bivalent Booster. N Engl J Med 388, 567–569. 10.1056/NEJMc2213907.

4. Wang, Q., Bowen, A., Tam, A.R., Valdez, R., Stoneman, E., Mellis, I.A., Gordon, A., Liu, L., and Ho, D.D. (2023). SARS-CoV-2 neutralising antibodies after bivalent versus monovalent booster. Lancet Infect Dis 23, 527–528. 10.1016/S1473-3099(23)00181-0.

5. Wang, Q., Guo, Y., Tam, A.R., Valdez, R., Gordon, A., Liu, L., and Ho, D.D. (2023). Deep immunological imprinting due to the ancestral spike in the current bivalent COVID-19 vaccine. Cell Rep Med, 101258. 10.1016/j.xcrm.2023.101258.

6. Wang, Q., Bowen, A., Ho, J., Zhang, R.M., Valdez, R., Stoneman, E., Gordon, A., Liu, L., and Ho, D.D. (2023). SARS-CoV-2 neutralising antibodies after a second BA.5 bivalent booster. Lancet 402, 1827–1828. 10.1016/S0140-6736(23)02278-X.

7. Collier, A.Y., Miller, J., Hachmann, N.P., McMahan, K., Liu, J., Bondzie, E.A., Gallup, L., Rowe, M., Schonberg, E., Thai, S., et al. (2023). Immunogenicity of BA.5 Bivalent mRNA Vaccine Boosters. N Engl J Med 388, 565–567. 10.1056/NEJMc2213948.

8. Kosugi, Y., Kaku, Y., Hinay, A.A., Jr., Guo, Z., Uriu, K., Kihara, M., Saito, F., Uwamino, Y., Kuramochi, J., Shirakawa, K., et al. (2024). Antiviral humoral immunity against SARS-CoV-2 omicron subvariants induced by XBB.1.5 monovalent vaccine in infection-naive and XBB-infected individuals. Lancet Infect Dis. 10.1016/S1473-3099(23)00784-3.

9. Patel, N., Trost, J.F., Guebre-Xabier, M., Zhou, H., Norton, J., Jiang, D., Cai, Z., Zhu, M., Marchese, A.M., Greene, A.M., et al. (2023). XBB.1.5 spike protein COVID-19 vaccine induces broadly neutralizing and cellular immune responses against EG.5.1 and emerging XBB variants. Sci Rep 13, 19176. 10.1038/s41598-023-46025-y.

10. Stankov, M.V., Hoffmann, M., Gutierrez Jauregui, R., Cossmann, A., Morillas Ramos, G., Graalmann, T., Winter, E.J., Friedrichsen, M., Ravens, I., Ilievska, T., et al. (2024). Humoral and cellular immune responses following BNT162b2 XBB.1.5 vaccination. Lancet Infect Dis 24, e1–e3. 10.1016/S1473-3099(23)00690-4.

11. Jain, S., Kumar, S., Lai, L., Linderman, S., Malik, A.A., Ellis, M.L., Godbole, S., Solis, D., Sahoo, M.K., Bechnak, K., et al. (2024). XBB.1.5 monovalent booster improves antibody binding and neutralization against emerging SARS-CoV-2 Omicron variants. bioRxiv, 2024.2002.2003.578771. 10.1101/2024.02.03.578771.

12. Liang, C.-Y., Raju, S., Liu, Z., Li, Y., Arunkumar, G.A., Case, J.B., Zost, S.J., Acreman, C.M., Anjos, D.C.C.d., McLellan, J.S., et al. (2024). Prototype mRNA vaccines imprint broadly neutralizing human serum antibodies after Omicron variant-matched boosting. bioRxiv, 2024.2001.2003.574018. 10.1101/2024.01.03.574018.

13. Chalkias, S., McGhee, N., Whatley, J.L., Essink, B., Brosz, A., Tomassini, J.E., Girard, B., Wu, K., Edwards, D.K., Nasir, A., et al. (2023). Safety and Immunogenicity of XBB.1.5-Containing mRNA Vaccines. medRxiv, 2023.2008.2022.23293434. 10.1101/2023.08.22.23293434.

14. Modjarrad, K., Che, Y., Chen, W., Wu, H., Cadima, C.I., Muik, A., Maddur, M.S., Tompkins, K.R., Martinez, L.T., Cai, H., et al. (2023). Preclinical Characterization of the Omicron XBB.1.5-Adapted BNT162b2 COVID-19 Vaccine. bioRxiv, 2023.2011.2017.567633. 10.1101/2023.11.17.567633.

15. Marking, U., Bladh, O., Aguilera, K., Yang, Y., Greilert Norin, N., Blom, K., Hober, S., Klingstrom, J., Havervall, S., Aberg, M., et al. (2024). Humoral immune responses to the monovalent XBB.1.5-adapted BNT162b2 mRNA booster in Sweden. Lancet Infect Dis 24, e80–e81. 10.1016/S1473-3099(23)00779-X.

16. Link-Gelles, R., Ciesla, A.A., Mak, J., Miller, J.D., Silk, B.J., Lambrou, A.S., Paden, C.R., Shirk, P., Britton, A., Smith, Z.R., and Fleming-Dutra, K.E. (2024). Early Estimates of Updated 2023-2024 (Monovalent XBB.1.5) COVID-19 Vaccine Effectiveness Against Symptomatic SARS-CoV-2 Infection Attributable to Co-Circulating Omicron Variants Among Immunocompetent Adults - Increasing Community Access to Testing Program, United States, September 2023-January 2024. MMWR Morb Mortal Wkly Rep 73, 77–83. 10.15585/mmwr.mm7304a2.

17. Simon, V., Kota, V., Bloomquist, R.F., Hanley, H.B., Forgacs, D., Pahwa, S., Pallikkuth, S., Miller, L.G., Schaenman, J., Yeaman, M.R., et al. (2022). PARIS and SPARTA: Finding the Achilles’ Heel of SARS-CoV-2. mSphere 7, e0017922. 10.1128/msphere.00179-22.

18. Wang, Q., Guo, Y., Iketani, S., Nair, M.S., Li, Z., Mohri, H., Wang, M., Yu, J., Bowen, A.D., Chang, J.Y., et al. (2022). Antibody evasion by SARS-CoV-2 Omicron subvariants BA.2.12.1, BA.4 and BA.5. Nature 608, 603–608. 10.1038/s41586-022-05053-w.

19. Smith, D.J., Lapedes, A.S., de Jong, J.C., Bestebroer, T.M., Rimmelzwaan, G.F., Osterhaus, A.D., and Fouchier, R.A. (2004). Mapping the antigenic and genetic evolution of influenza virus. Science 305, 371–376. 10.1126/science.1097211.

